# A *C. elegans* neuron both promotes and suppresses motor behavior to fine tune motor output

**DOI:** 10.1101/2020.11.02.354472

**Authors:** Zhaoyu Li, Jiejun Zhou, Khursheed Wani, Teng Yu, Elizabeth A. Ronan, Beverley J. Piggott, Jianfeng Liu, X.Z. Shawn Xu

**Affiliations:** Queensland Brain Institute, University of Queensland, St Lucia, QLD 4072, Australia; Life Sciences Institute and Department of Molecular and Integrative Physiology, University of Michigan, Ann Arbor, MI 48109, USA; College of Life Science and Technology, Key laboratory of Molecular Biophysics of MOE, and International Research Center for Sensory Biology and Technology of MOST, Huazhong University of Science and Technology, Wuhan, Hubei, 430074, China; Department of Microbiology and Physiological Systems, University of Massachusetts Medical School, Worcester, MA 01605, USA; Division of Biological Sciences, University of Montana, Missoula, MT 59812

## Abstract

How neural circuits drive behavior is a central question in neuroscience. Proper execution of motor behavior requires the precise coordination of many neurons. Within a motor circuit, individual neurons tend to play discrete roles by promoting or suppressing motor output. How exactly neurons function in specific roles to fine tune motor output is not well understood. In *C. elegans*, the interneuron RIM plays important yet complex roles in locomotion behavior. Here, we show that RIM both promotes and suppresses distinct features of locomotion behavior to fine tune motor output. This dual function is achieved via the excitation and inhibition of the same motor circuit by electrical and chemical neurotransmission, respectively. Additionally, this bi-directional regulation contributes to motor adaptation in animals placed in novel environments. Our findings reveal that individual neurons within a neural circuit may act in opposing ways to regulate circuit dynamics to fine tune behavioral output.

## Introduction

Animals execute a wide range of behaviors, which rely on the vast number of neurons in the brain. Control of motor output is an essential feature of the nervous system in nearly all animals, and the successful execution of even the simplest motor behaviors requires the precise coordination of many individual neurons (Purves et al., 2008). For example, a simple withdrawal behavior in land snails involves different groups of neurons including sensory, motor, modulatory, and command neurons (Balaban, 2002). Individual neurons within circuits tend to play discrete roles in either promoting or suppressing motor output. For example in the mammalian motor cortex output circuit, distinct neurons release glutamate or GABA, to form a feedforward excitatory or inhibitory circuit, respectively, to regulate motor outputs (Cote et al., 2018). However, how individual neurons coordinate within a functional circuit to generate motor output is not well understood.

*C. elegans* has emerged as a highly valuable model to investigate the mechanisms by which neural circuits control behavior. *C. elegans* possess a simple nervous system composed of 302 neurons, approximately 7000 chemical synapses, and 900 electrical junctions (White et al., 1986). These elements together generate a wide variety of behaviors, ranging from simple behaviors such as sensory detection and motor output to more complex behaviors including mating, sleep, drug-dependency, and learning (de Bono and Maricq, 2005; Feng et al., 2006; Hart and Chao, 2010; Pierce-Shimomura et al., 2008). Furthermore, the connectome of *C. elegans* nervous system has been mapped in exquisite detail by electron microscopy reconstruction, although this only reveals structural but not functional connections. These features together make *C. elegans* an excellent model to investigate the neural and genetic mechanisms by which individual neurons function within a circuit to drive motor output.

In order to navigate the environment, *C. elegans* locomotion is driven by undulations propagating from head to tail. Reorientation via backward locomotion, also called reversal, is a key behavioral strategy in animal navigation and avoidance of aversive stimuli (Gray et al., 2005; Hilliard et al., 2002; Piggott et al., 2011). Despite its simplicity, several elements of this motor program must be elaborately controlled to ensure it proper execution. This includes regulation of timing and strength of the motor output, as well as the likelihood that the behavior is initiated in a specific instance, termed response probability. Many neurons are involved in reversal regulation, ranging from the most upstream sensory neurons down to motor neurons (Gray et al., 2005). Laser ablation studies showed that the interneurons AVA and AVE command reversal execution through the A-type motor neurons (Chalfie et al., 1985). This is further corroborated by calcium imaging studies revealing that the activities of AVA/AVE neurons are tightly coupled with reversals (Kato et al., 2015; Kawano et al., 2011; Piggott et al., 2011). The command interneurons AVA/AVE form a large number of connections with the first layer and second layer interneurons, which are thought to relay sensory information (White et al., 1986). While AVA/AVE command interneurons are essential drivers of reorientation during locomotion, less is understood regarding exactly how these neurons are regulated within the locomotion circuitry to control motor output.

Among the second layer interneurons that connect with AVA/AVE, many reports implicate the pair of RIM interneurons as having an important role in reversal regulation. RIM neurons form both electrical and chemical synapses with AVA/AVE neurons. Laser ablation of RIM neurons has been reported to increase the frequency of reversals, suggesting an inhibitory role of RIM neurons in reversal regulation (Gray et al., 2005; Piggott et al., 2011). Interestingly, RIM-ablated worms also exhibit a reduction in reversal responses to anterior tactile stimulation or osmolarity insult, indicating a promotion role of RIM neurons in reversal regulation (Piggott et al., 2011; Zheng et al., 1999). Furthermore, the calcium activity in RIM is coupled with reversals (Kato et al., 2015; Kawano et al., 2011). While these findings highlight the important role of RIM in regulating reversal behavior, they also reveal a critical knowledge gap in our understanding of how RIM functions in the locomotion circuitry to drive reversal behavior.

In the present study, we investigated how RIM functions and coordinates with AVA/AVE command interneurons to form a functional circuit that properly controls reversal behavior. By combining optogenetics, laser ablation, calcium imaging and molecular genetics, we interrogated the complex roles of RIM in regulating distinct features of reversal behavior. We found that while RIM functions to promote reversal with AVA/AVE, it also suppresses reversal via AVA/AVE and A-type motor neurons. At the molecular level, RIM’s promotion of reversal behavior requires gap junctions with AVA/AVE, while its suppression role relies on chemical neurotransmission with AVA/AVE and A-type motor neurons. At the circuit level, RIM can both promote and suppress AVA/AVE neuronal activities. Additionally, we uncovered that this bi-directional regulation of neural circuits is involved not only in the simple reversal behavior, but also in more complex behaviors such as motor adaptation. Our work identifies circuit and molecular mechanisms by which individual neurons within a neural circuit both promote and suppress motor behavior to fine tune motor output.

## Results

### The pair of RIM interneurons promote reversal initiation while suppressing reversal probability

*C. elegans* locomotion consists of forward crawling interrupted with reversals. Reversal allows worms to change locomotion direction, a key behavioral strategy in navigation and avoidance of aversive stimuli (Gray et al., 2005). Previous reports indicate a complex role of RIM neurons in the regulation of reversal behavior. To investigate the role of RIM neurons in this behavior, we adopted two behavioral assays. Specifically, we employed optogenetics to evoke reversals acutely to assay reversal initiation. To assay reversal probability, we recorded the frequency of spontaneous reversal events during worm locomotion.

AVA and AVE neurons act as command interneurons for reversal behavior (Chalfie et al., 1985; Gray et al., 2005; Piggott et al., 2011). Indeed, acute activation of AVA neurons optogenetically with Chrimson, a red light-drivable channelrhodopsin (Klapoetke et al., 2014), triggered reversals immediately (Figure 1A-B), confirming the positive role of AVA neurons in reversal initiation. A similar phenomenon was observed with AVE neurons (Figure 1C-D). When AVA and AVE neurons were ablated, the reversal frequency was greatly reduced (Figure 1G), verifying a critical role of AVA/AVE in promoting reversal probability. Notably, in AVA/AVE-ablated worms, the length (head swings) of the residual reversal events was rather short (Figure 1H). These results support the notion that AVA and AVE neurons play a critical role in driving reversal behavior.

**Figure 1.**
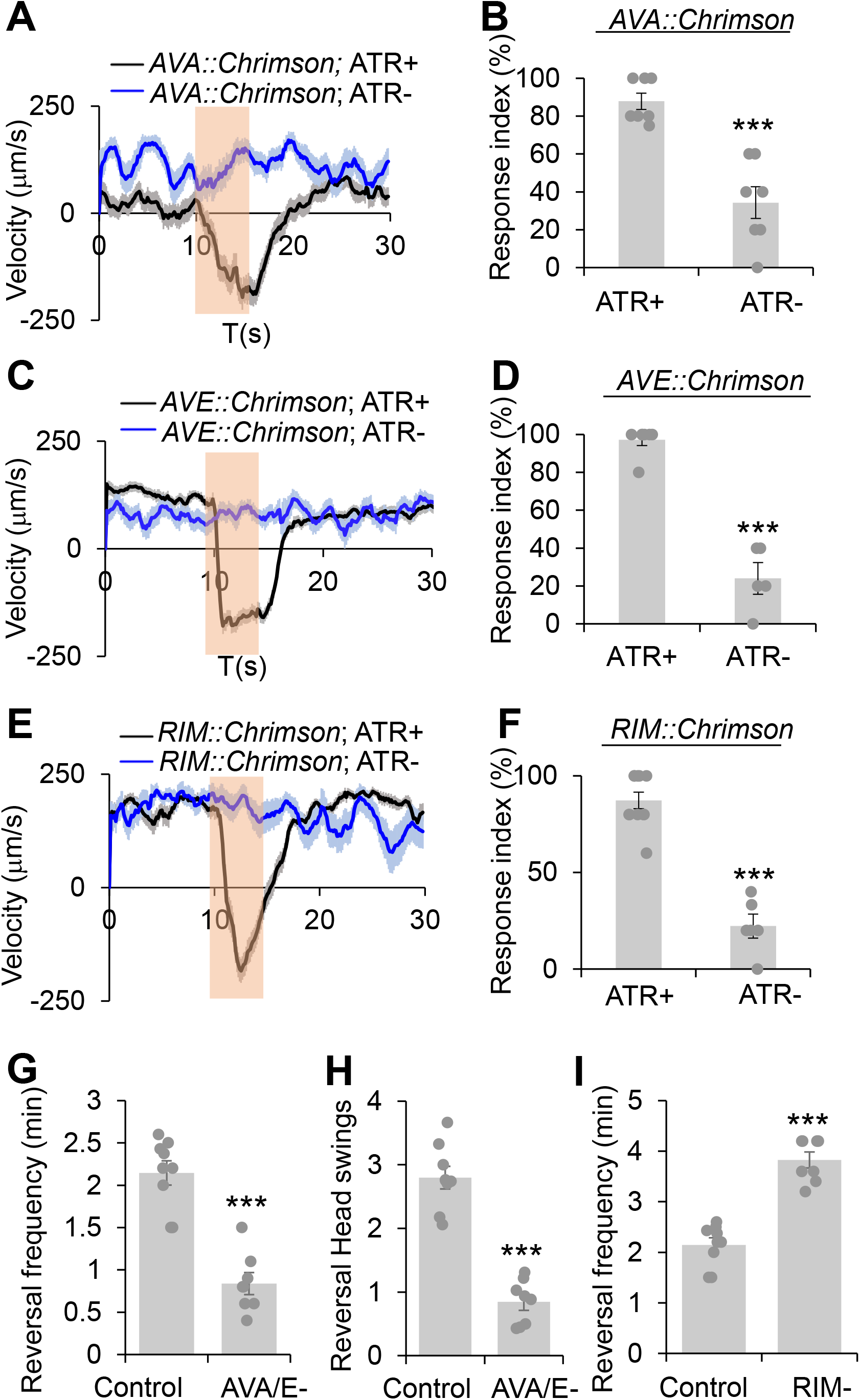
RIM has a complex role in reversal regulation. **(A-B)** Acute activation of AVA neurons using Chrimson triggers reversals. AVA was stimulated optogenetically by a Chrimson transgene under the *npr-4* promoter. (A) Average velocity trace with SEM. n>=35. **(B)** Bar graph of reversal index quantification. Error bars: SEM. n≥7. ***p=0.000106 (unpaired two-sided t-test). ATR: all-trans retinal, which is required for the function of Chrimson. The bar in amber denotes the time window of light illumination. **(C-D)** Acute activation of AVE neurons using Chrimson trigger reversals. AVE was stimulated optogenetically by a Chrimson transgene under the *opt-3* promoter. (C) Average velocity trace with SEM. n>=25. (D) Bar graph of reversal index quantification. Error bars: SEM. n≥5. ***p=1.18e-6 (unpaired two-sided t-test). **(E-F)** Acute activation of RIM neurons using Chrimson triggers reversals. RIM was stimulated optogenetically by a Chrimson transgene under the *gcy-13* promoter. (E) Average velocity trace with SEM. n>=30. F. Bar graph of reversal index quantification. Error bars: SEM. n≥6. ***p=1.063e-7 (unpaired two-sided t-test). **(G-H)** Ablation of AVA and AVE neurons reduces reversal frequency and reversal head swings. (G) Quantification of reversal frequency. Error bars: SEM. n≥8. ***p= 3.529e-6 (unpaired two-sided t-test). Quantification of reversal headswings. (H) Error bars: SEM. n≥8. ***p=4.919e-7 (unpaired two-sided t-test). **(I**) Ablation of RIM neurons increases reversal frequency. Bar graph shows average reversal frequency. Error bars: SEM. n≥8. ***p=2.39e-5 (unpaired two-sided t-test).

RIM neurons form both gap junctions and chemical synapses with AVA and AVE neurons (White et al., 1986). To test if RIM neurons share roles similar to AVA/AVE neurons in reversal regulation, we conducted both optogenetic and laser ablation experiments. Similar to AVA/AVE neurons, optogenetic activation of RIM neurons using Chrimson rapidly triggered reversals (Figure 1E-F), confirming a role for RIM neurons in promoting reversal initiation (Guo et al., 2009; Zheng et al., 1999). On the other hand, as reported previously (Gray et al., 2005; Piggott et al., 2011), ablation of RIM resulted in an increase in reversal frequency, indicating a role of RIM neurons in suppressing reversal probability (Figure 1I). Thus, RIM neurons appear to both promote and suppress reversal behavior.

To further interrogate the function of AVA/AVE and RIM neurons in reversal regulation, we recorded the calcium activities of these neurons in freely-behaving worms with the genetic calcium indicator GCaMP6 (Chen et al., 2013), using an automated calcium imaging system. We observed that reversal events were tightly coupled with the rising phase of calcium spikes in AVA/AVE and RIM neurons, supporting the idea that both neurons contribute to reversal initiation (Figure S1A-B). Altogether, the above results support the notion that while the command interneurons AVA/AVE are important in promoting reversal, the interneuron RIM plays a more complex role by promoting reversal initiation and suppressing reversal probability.

### RIM neurons promote reversal initiation via gap junctions

Having shown that RIM neurons possess roles in both the promotion of reversal initiation and suppression of reversal probability, we next asked how this dual-function is achieved at the circuit and molecular levels. The *C. elegans* wiring diagram reveals that RIM forms both chemical synapses and electrical gap junctions with AVA and AVE. As AVA/AVE are known to mediate reversal behavior (Chalfie et al., 1985; White et al., 1986), we sought to determine whether RIM promotes reversal initiation through these neurons. A short pulse of red light rapidly triggered a reversal in *RIM::Chrimson* worms (Figure 2A-B). However, when AVA and AVE neurons were removed by laser ablation, red light was no longer able to trigger reversals in *RIM::Chrimson* worms, although a decrease in forward speed was still observed (Figure 2A-B). These results suggest that the command interneurons AVA/AVE are required for RIM to promote reversal initiation.

**Figure 2.**
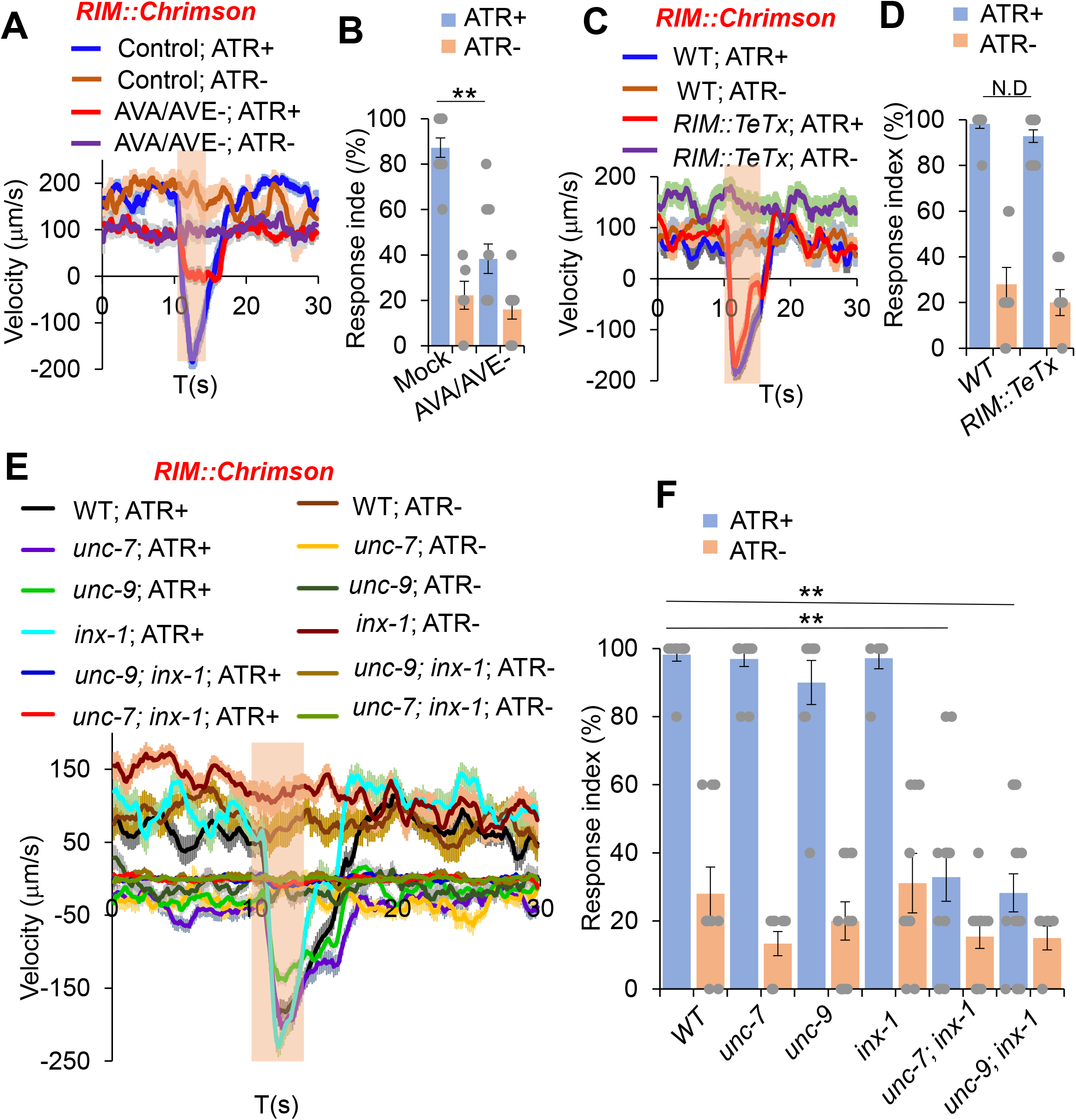
RIM promotes reversals through command interneurons and gap junctions. **(A-B)** Ablation of AVA/AVE neurons decreases RIM::Chrimson-triggered reversals. RIM was stimulated optogenetically with a Chrimson transgene under the *gcy-13* promoter. (A) Traces shows average speed with SEM. n>=30. ATR: all-trans retinal. The bar in amber denotes the time window of light illumination. (B) Reversal index quantification of (A). Error bars: SEM. n≥6. **p=1.095e-05 (ANOVA with Tukey’s HSD test). **(C-D)** Blockade of chemical transmission using Tetanus toxin (TeTx) in RIM does not change RIM::Chrimson-triggered reversals. TeTx was expressed in RIM as a transgene using the *gcy-13* promoter. (C) Average velocity traces with SEM. n>=40. (D) Reversal index quantification of (C). Error bars: SEM. n≥8. p=0.7992 (ANOVA with Tukey’s HSD test). **(E-F)** Gap junction mutants impair RIM::Chrimson-triggered reversals. (E) Average velocity traces with SEM. n>=40. (F) Reversal index quantification of (E). Error bars: SEM. n≥8. (*unc-7;inx-1*, p=1.048e-05; *unc-9;inx-1*, p=1.048e-05 (ANOVA with Tukey’s HSD test)).

As RIM forms both chemical and electrical synapses with AVA and AVE neurons (Chalfie et al., 1985; White et al., 1986), we asked which type of synapses mediate the transmission between RIM and AVA/AVE. To test whether chemical synapses are required, we employed genetically coded toxins to block chemical synapses. Tetanus toxin (TeTx) specifically cleaves synaptobrevin to impair chemical neurotransmission (Pellizzari et al., 1999). We expressed TeTx as a transgene in RIM using a RIM-specific promoter, and tested whether *RIM::Chrimson*-triggered reversals were affected. Impairment of RIM chemical transmission with TeTx did not block *RIM::Chrimson*-triggered reversals (Figure 2C-D), suggesting that chemical synapses are not required for RIM neurons to promote reversal initiation through AVA/AVE neurons.

We next examined whether electrical synapses play a role in RIM-triggered reversal initiation. Previous work reported that two innexin genes *unc-7* and *unc-9* are expressed in RIM and AVA/AVE neurons (Altun et al., 2009; Bhattacharya et al., 2019). We found that another innexin gene, *inx-1,* was also highly expressed in these neurons (Figure S2). To test whether these innexins mediate the transmission between RIM and AVA/AVE neurons, we activated RIM neurons in *unc-7*, *unc-9* and *inx-1* single or double mutant animals using Chrimson. None of the single gap junction mutants showed a defect in *RIM::Chrimson*-triggered reversals (Figure 2E-F). However, *unc-7; inx-1* and *unc-9; inx-1* double mutants exhibited largely reduced responses (Figure 2E-F), suggesting that these innexins function in combination to mediate electrical transmission between RIM and AVA/AVE neurons. It should be noted that these double mutants displayed a more severe uncoordinated phenotype than *unc-7* and *unc-9* single mutants. To ensure that the reduced responses in innexin double mutants were not simply caused by uncoordinated movement, we optogenetically activated the downstream command neurons AVA in *unc-7; inx-1* double mutant worms. We observed that upon activation of AVA neurons, *unc-7; inx-1* worms were still able to execute reversals, albeit at a reduced speed and response rate (Figure S3A-B). Thus, these double mutants retained the ability to execute reversals, though they failed to do so upon activation of RIM. These results demonstrate that the gap junction genes *inx-1, unc-7* and *unc-9* contribute to RIM-triggered reversal initiation. We thus conclude that RIM promotes reversal initiation via AVA/AVE command interneurons through gap junctions.

### RIM neurons suppress reversal probability via chemical neurotransmission

We next asked how RIM suppresses reversal probability. Previous findings indicate that loss of the first layer interneurons AIB and AIZ, the command interneurons AVA and AVE, and the A-type motor neurons decreases the reversal frequency (Chalfie et al., 1985; Gray et al., 2005; Li et al., 2014). To test whether any of these neurons function downstream of RIM to mediate its suppression effect on reversal probability, we removed these neurons by laser ablation and tested if it eliminated the hyper-reversal phenotype caused by the loss of RIM neurons.

Removal of AIB and AIZ neurons did not abolish the hyper-reversal phenotype in RIM-ablated animals, suggesting that AIB and AIZ neurons do not function downstream of RIM to suppress reversal probability (Figure S4A). In contrast, when the AVA/AVE or A-type motor (Amo) neurons were removed by laser ablation or functionally impaired by TeTx, the hyper-reversal phenotype in RIM-ablated worms was largely suppressed, suggesting that RIM suppresses reversal probability via the AVA/AVE-Amo circuit (Figure S4B-C).

Having characterized the circuit mechanism by which RIM suppresses reversals, we next sought to identify the underlying molecular mechanisms. We again asked whether chemical synaptic transmissions were required. To address this, we specifically expressed TeTx in RIM neurons as a transgene to block their chemical transmission, and recorded the spontaneous reversal frequency. We found that blocking chemical transmission in RIM neurons increased reversal frequency, suggesting that RIM suppresses reversal probability via chemical transmission (Figure 3A).

**Figure 3.**
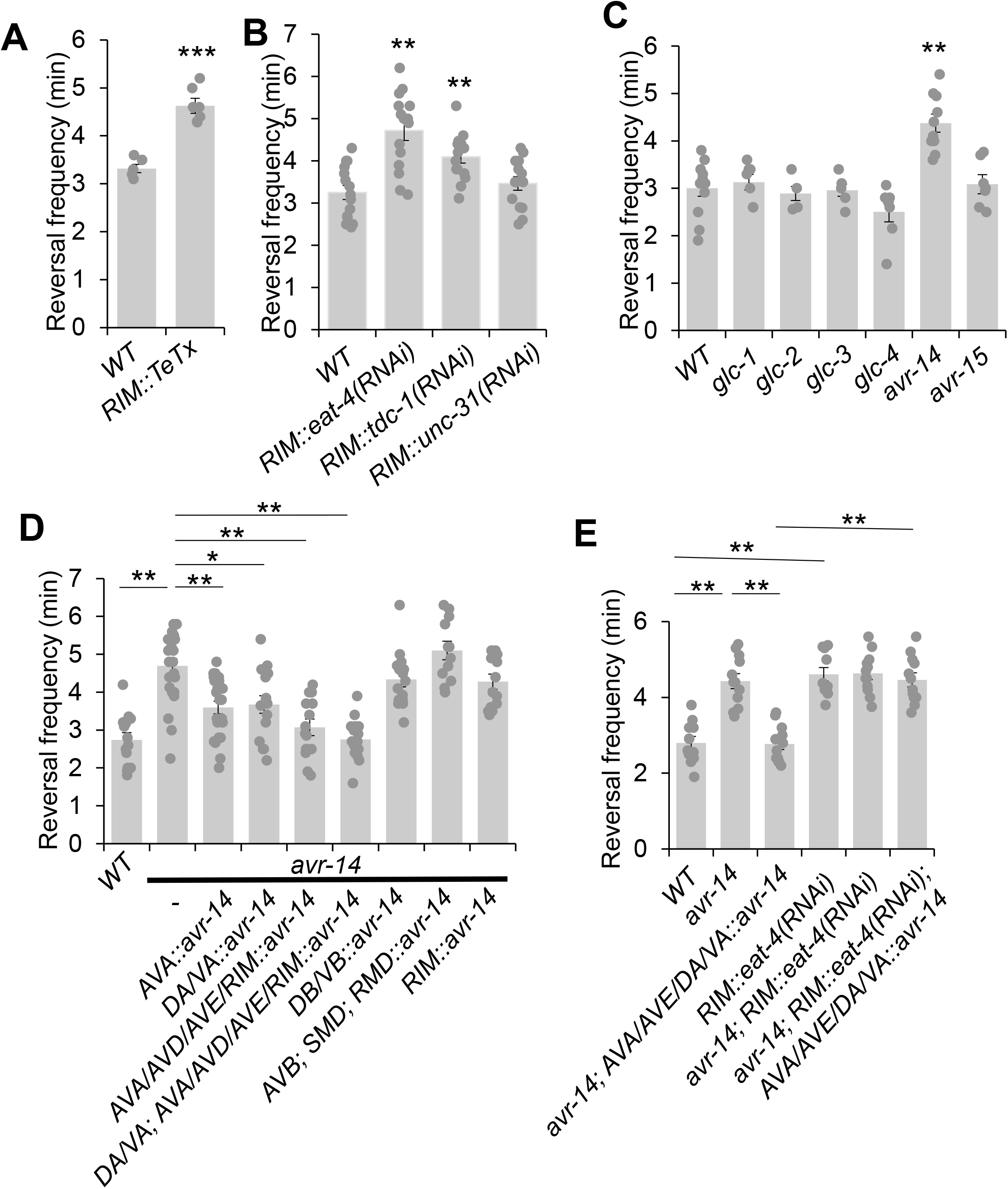
RIM suppresses reversal frequency by chemical transmission. **(A)** Blockage of RIM chemical transmission using Tetanus toxin (TeTx) in RIM increases reversal frequency. Bar graph shows quantification of reversal frequency. Error bars: SEM. n≥6. *** p=2.73e-05 (unpaired two-sided t-test). **(B)** Knocking down glutamate and tyramine release by RNAi in RIM increases reversal frequency. RNAi was expressed as a transgene in RIM using the *gcy-13* promoter. Bar graph shows quantification of reversal frequency. Error bars: SEM. n≥14. *RIM::eat-4(RNAi)*, **p=1.171e-05. *RIM::tdc-1(RNAi),* **p=0.006846 (ANOVA with Tukey’s HSD test). **(C)** Mutation in *avr-14* but not other glutamate-gated chloride channel genes increases the reversal frequency. Bar graph shows quantification of reversal frequency. Error bars: SEM. n≥4. **p=0.0003544 (ANOVA with Tukey’s HSD test). **(D)** Transgenic expression of wild-type *avr-14* gene in AVA/AVE command interneurons and A-type motor neurons rescues the hyper-reversal defect of *avr-14* mutant worms. Bar graph shows quantification of reversal frequency. Error bars: SEM. n≥14. **p=9.89e-06 between WT and *avr-14;* **p= 0.001245 between *avr-14* and *AVA::avr-14 rescue;* *p= 0.01786 between *avr-14* and *DA/VA::avr-14* rescue; **p=1.324e-05 between *avr-14* and *AVA/AVD/AVE/RIM::avr-14* rescue; **p=9.887e-06 between *avr-14* and *DA/VA; AVA/AVD/AVE/RIM::avr-14* rescue (ANOVA with Tukey’s HSD test). **(E)** Blockage of glutamate release from RIM by *RIM::eat-4(RNAi)* impairs the rescue of *avr-14* phenotype mediated by an *avr-14* transgene expressed in in AVA/AVE and A-type motor neurons. Bar graph shows quantification of reversal frequency. Error bars: SEM. n≥10. **p=1.022e-05 between WT and *avr-14; \*\**p=1.018e-05 between WT and *RIM::eat-4(RNAi). \*\**p= 1.019e-05 between *avr-14* and *avr-14; AVA/AVE/DA/VA::avr-14. \*\**p=1.018e-05 between *avr-14; RIM::eat-4(RNAi); AVA/AVE/DA/VA::avr-14* and *avr-14; AVA/AVE/DA/VA::avr-14*) (ANOVA with Tukey’s HSD test). Promoters that drive *avr-14* transgene expression are *Pnpr-4*: AVA neurons; *Punc-4*: DA, VA neurons; *Pnmr-1*: AVA, AVE, RIM and AVD; *Pacr-5*: DB, VB motor neurons; *Plgc-55*: AVB, SMD, RMD neurons; *Pgcy-13*: RIM neurons.

Chemical transmission in the nervous system is typically mediated by classic neurotransmitters and neuropeptides. RIM neurons release glutamate, tyramine and possibly neuropeptides (Alkema et al., 2005; Kim and Li, 2004; Serrano-Saiz et al., 2013). To test which of these is required for RIM’s suppression effect on reversal probability, we specifically knocked down the associated pathways in RIM with RNAi of the following genes: *eat-4*, which encodes a vesicle glutamate transporter (Lee et al., 1999); *tdc-1* which encodes a tyrosine decarboxylase required for tyramine biogenesis (Alkema et al., 2005); and *unc-31*, which is required for neuropeptide release (Speese et al., 2007). No effect was observed in *unc-31(RNAi)* worms, suggesting that neuropeptide signaling may not play a major role in mediating the suppression effect of RIM on reversal probability (Figure 3B). By contrast, knocking down glutamate release from RIM markedly increased the reversal frequency, suggesting that glutamate release from RIM may suppress reversal probability (Figure 3B). A similar result was obtained with *tdc-1* knock down, although the effect was not as robust as that observed with *eat-4* knockdown (Figure 3B). Tyramine is known to suppress reversal frequency via tyramine-gated chloride channels (Pirri et al., 2009). However, how glutamatergic signaling suppresses reversal frequency is completely unknown. Given this and the fact that RNAi of *eat-4* exhibited a more robust effect, we focused on characterizing the role of glutamate transmission in the suppression of reversal probability.

To identify the glutamate receptor that acts downstream of RIM neurons to mediate the suppression of reversal probability, we examined various glutamate receptor mutants. We focused on glutamate-gated chloride channels as they are known to mediate inhibitory responses (Dent et al., 2000). Mutant worms lacking *avr-14*, which encodes a glutamate-gated chloride channel (Dent et al., 2000), exhibited a hyper-reversal phenotype, similar to that detected in RIM-ablated and *RIM::eat-4(RNAi)* worms (Figure 3C). In addition, we found that *avr-14* was expressed in neurons including AVA/AVE and A-type motor neurons (Figure S4D). Furthermore, transgenic expression of wild-type *avr-14* gene in AVA/AVE and A-type motor (Amo) neurons rescued the hyper-reversal phenotype in *avr-14* mutant worms (Figure 3D). Moreover, blocking glutamate release specifically from RIM by *RIM::eat-4(RNAi)* abolished the rescue effect of the *avr-14* transgene (Figure 3E). Taken together, these results suggest that the glutamate-gated chloride channel AVR-14 functions in the AVA/AVE-Amo circuit to mediate the suppression of reversal probability by RIM neurons.

### RIM both promotes and suppresses AVA/AVE activities

Our results show that RIM neurons can both promote reversal initiation as well as suppresses reversal probability through interactions with the interneurons AVA/AVE. We next wondered how RIM regulates the activities of AVA/AVE neurons. To address this, we examined how RIM ablation affects the calcium activities in AVA/AVE neurons in freely-moving worms. In mock-ablated controls, the calcium spikes in AVA/AVE neurons were coupled with reversals, with reversals initiating upon calcium increase in AVA/AVE, and terminating once calcium traces peaked and began to drop (Figure 4A, Figure S5B, D). In RIM-ablated animals, we observed that the coupling between reversal events and calcium spikes as well as the kinetics of calcium spikes were still preserved (Figure S5A-D). However, the amplitude of AVA/AVE calcium spikes was significantly reduced in RIM-ablated worms (Figure 4A-C). These results suggest that RIM promotes AVA/AVE neuronal activities by increasing the amplitude of individual calcium spikes without changing their kinetics. This is further supported by the amplitude distribution pattern of individual calcium spikes (Figure S5A). Specifically, in RIM-ablated animals, the amplitude distribution pattern is left-shifted to a narrower window, indicating that the calcium spikes in AVA/AVE neurons became weaker in the absence of RIM neurons (Figure S5A).

**Figure 4.**
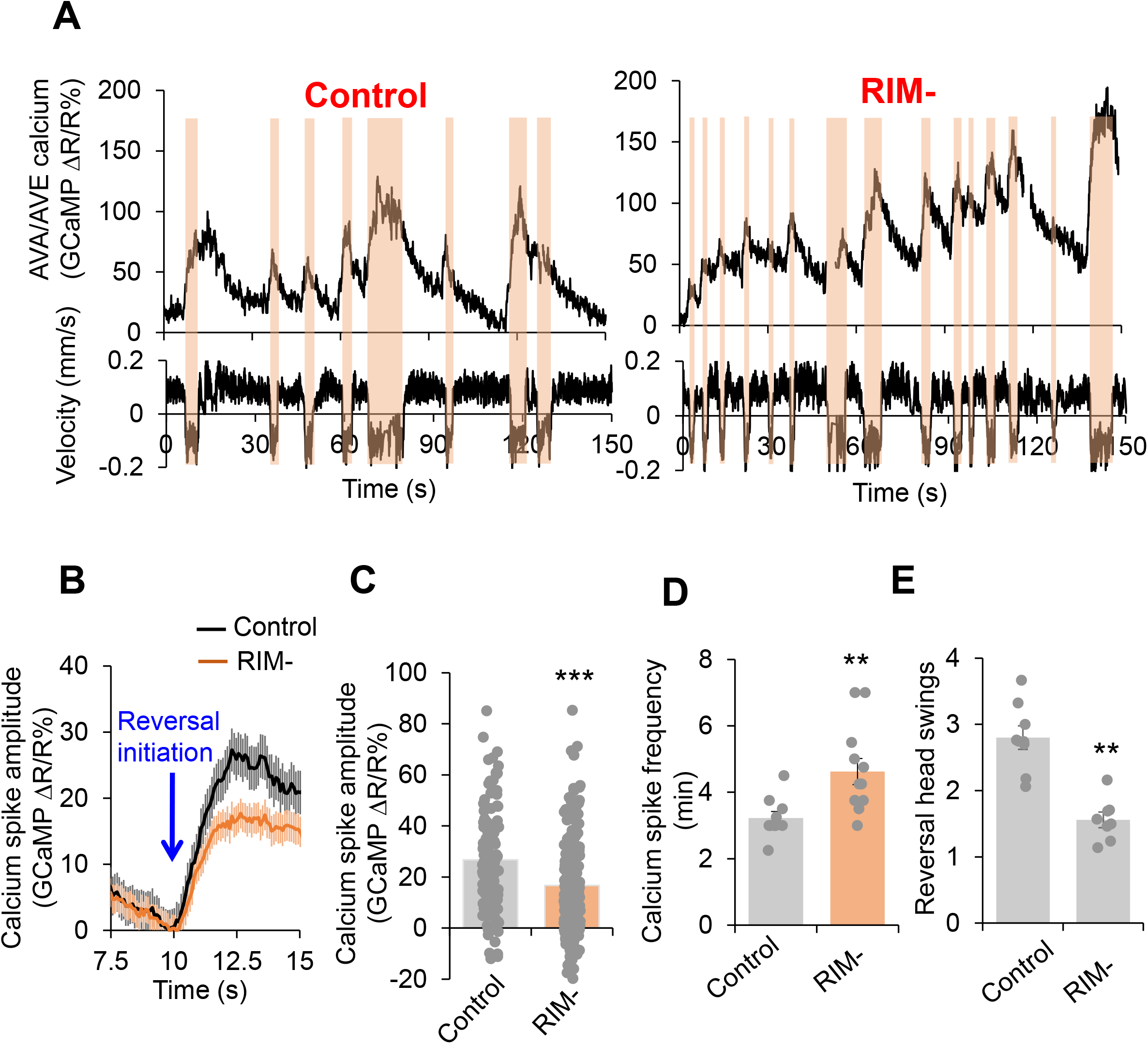
RIM fine tunes AVA/AVE activities. **(A)** Calcium spikes in AVA/AVE neurons in freely-moving worms are tighly coupled with reversals. Calcium imaging was conducted with freely moving animals using the CARIBN system. Left panels: mock-ablated worms. Right panels: RIM-ablated worms. Upper panels: calcium traces. Lower panels: velocity traces. Amber bars label reversal events. The *nmr-1* promoter was used to drive GCaMP3/DsRed expression as a transgene in AVA and AVE neurons. As AVA and AVE neurons are in close proximity, the detected calcium fluorescence signals reflect the overall calcium activity in both neurons, though the calcium signals should be mainly contributed by AVA neurons due to the much stronger expression of GCaMP in AVA than AVE. **(B-C)** RIM ablation decreases the amplitude of calcium spikes in AVA/AVE neurons. (B) Average traces with SEM. Blue arrow marks the time point of reversal initiation. (C) Bar graph shows quantification of the amplitude of calcium spikes. Error bars: SEM. n≥150. *** p=5.24e-8 (unpaired two-sided t-test). **(D)** RIM ablation increases the frequency of calcium spikes in AVA/AVE neurons. Bar graph shows quantification the frequency of calcium spikes. Error bars: SEM. n≥12. ** p=0.005311 (unpaired two-sided t-test). **(E)** Reversal length (head swings) is reduced in RIM ablated animals. Bar graph shows quantification of reversal head swings. Error bars: SEM. n≥8. ***p=2.39e-5 (unpaired two-sided t-test).

This calcium imaging result is consistent with the behavioral data in which we found the length of reversals (head swings) became shorter in RIM-ablated animals (Figure 4E). On the other hand, the frequency of calcium spikes in AVA/AVE neurons was increased in RIM-ablated worms (Figure 4A and 4D), indicating that AVA/AVE neurons became more excitable in the absence of RIM. Thus, RIM appears to both promote and suppress AVA/AVE activities. These findings also suggest that RIM promotes reversal initiation by potentiating the amplitude of calcium spikes in AVA/AVE neurons, but suppresses reversal probability by inhibiting the frequency of calcium spikes in these neurons, thereby providing a circuit mechanism underlying the dual-role of RIM in regulating reversal behavior.

### The dual-role of RIM neurons in motor adaptation

Given our observations that RIM can bi-directionally promote and suppress reversal behavior, we next wondered whether this dual function of RIM contributes to reversal-related complex behaviors under more natural conditions. One possible application of this function could be to facilitate motor adaptation after food removal. In the presence of food, worms execute mostly short reversals (less than one head swing) (Figure 5B). Upon transfer to a no-food environment, the reversal length markedly increased to >3 head swings (Figure 5B). Furthermore, the total reversal strength (reversal head swings multiplied by reversal frequency) increased dramatically in the first minute following transfer to a no-food environment (Figure 5C). Constantly maintaining such a high response is not beneficial to animals, as it would be energetically costly to sustain the behavior. Indeed, animals underwent fast motor adaptation following transfer to the no-food environment (Figure 5A-C).

**Figure 5.**
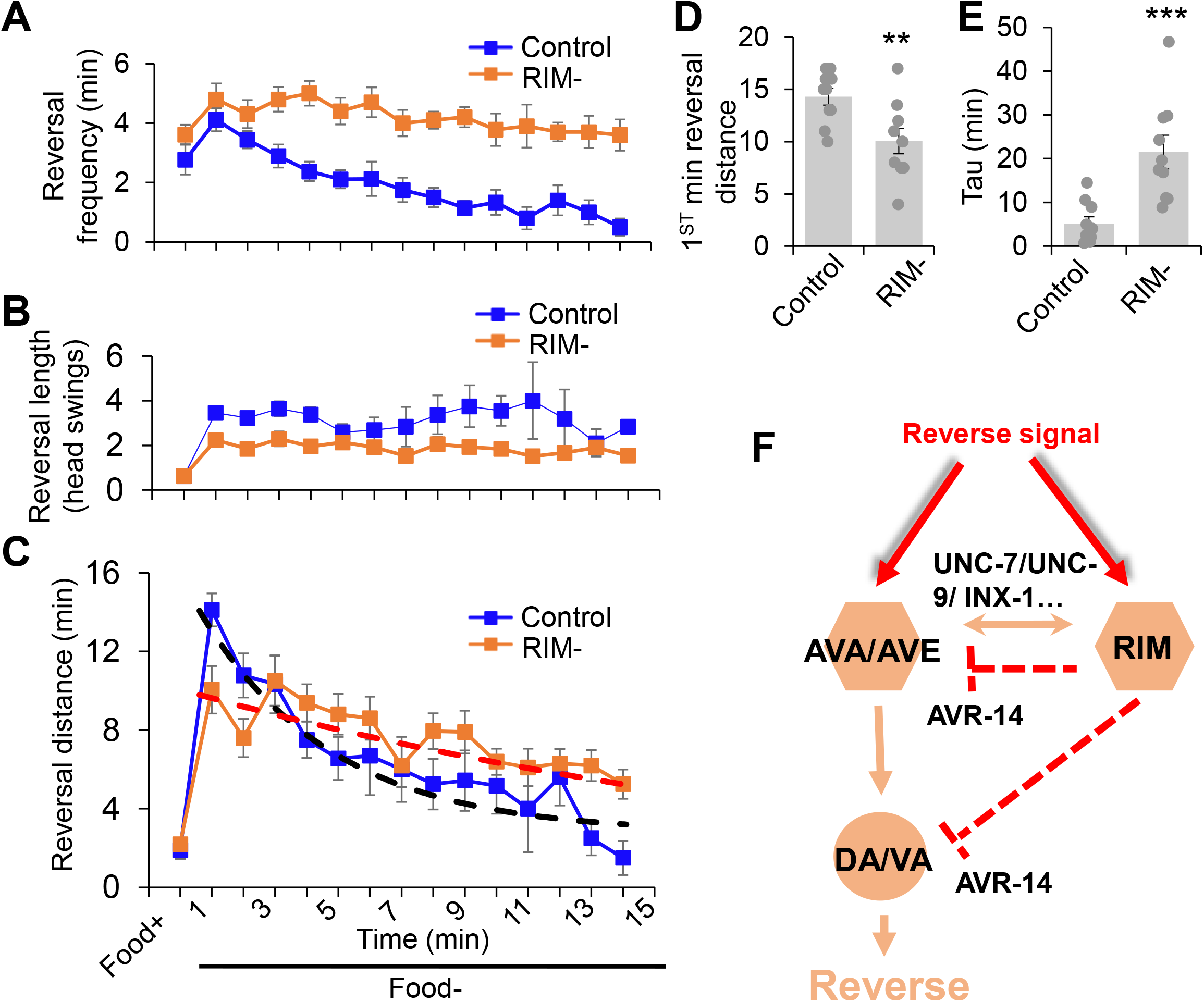
The dual role of RIM neurons in motor adaptation. Quantification of reversal properties in a motor adaptation assay. n=10. Control groups of animals underwent surgical preparation without laser irradiation. **(A)** RIM ablation blocks the reversal frequency decline after worms were transferred to no-food environment. Error bars: SEM. n=10. **(B)** RIM ablation decreases the reversal length after worms were transferred to no-food environment. SEM. n=10. **(C)** Ablation of RIM neurons impairs motor adaptation after worms were transferred to no-food environment. The reversal strength is the sum of the total reversal distance (reversal head swings) in each minute. Dash lines were the fitting curves for the reversal strength of control and RIM-ablated groups (fitted with exp function f(x)=y(0)+A*exp(-invTau*x). Control: y(0)=3.43, invTau=0.249, A=13.461; RIM-: y(0)=-0.07, invTau=0.042, A=10.467). SEM. n=10. **(D)** Ablation of RIM decreases the initial phase of the reversal strength (reversal strength of the 1st minute). Bar graph summarizes the 1st minute data in (C). **p=0.0066 (unpaired two-sided t-test). **(E)** Ablation of RIM led to a slower decline in the reversal strength indicated by Tau value. ***p=0.0006 (unpaired two-sided t-test). Tau values were derived from the fitting lines in (C). **(F)** Schematic model. RIM neurons acutely promotes reversals by promoting AVA/AVE activity via gap junctions. RIM also chronically inhibits AVA/AVE-A type motor neurons via an inhibitory glutamate pathway, thereby suppressing reversal probability over time.

Although the number of reversal head swings did not change over time in the no-food environment (Figure 5B), the reversal frequency quickly decreased over time (Figure 5A), resulting in a rapid drop in the total reversal strength over the 15 minute time window (Figure 5C), indicating fast motor adaptation. This adaptive behavior, also called local search behavior (Gray et al., 2005), featured two prominent phases: upon transfer to the no-food environment, worms first exhibited a rapid increase in reversal strength, followed by a progressive decrease in reversal strength over time (Figure 5C).

We then asked whether RIM neurons contribute to such motor adaptation. In RIM-ablated worms, the number of reversal head swings was decreased compared to controls following transfer to the no-food environment (Figure 5B), resulting in a significant decrease in total reversal strength in the initial phase (e.g. the first minute) of motor adaptation (Figure 5C and 5D). Despite this, as RIM-ablated worms displayed a much slower decline in the frequency of reversal events in the no-food environment (Figure 5A), the total reversal strength exceeded that observed in mock-ablated control worms in later phases of motor adaptation (e.g. >4 minute) (Figure 5C and 5E). The initial decrease in the reversal strength in RIM-ablated worms is consistent with RIM’s role in promoting reversal initiation, while the elevated reversal strength at later times is in line with RIM’s role in suppressing reversal probability. This biphasic defect in RIM-ablated worms supports the notion that RIM neurons both promote and suppress reversal behavior. Thus, RIM neurons contribute to motor adaptation in a new environment though their modulation of different features of reversal behavior. Taken together, our results provide a model in which RIM neurons function with AVA/AVE/A-type motor neurons to both promote and suppress the reversal circuit to fine tune motor output (Figure 5F).

## Discussion

Previous studies reported seemingly conflicting results with respect to the role of RIM neurons in the locomotion circuitry, suggesting an intricate role of RIM neurons in regulating locomotion. In the current study, we find that RIM neurons can both promote and suppress reversals during locomotion within a single motor circuit and do so by regulating distinct features of the reversal behavior. RIM neurons promote the initiation of individual reversal events while suppressing reversal probability. This multi-feature regulation is separately conducted by electrical and chemical transmissions with the reversal command interneurons AVA/AVE. We show that RIM promotes reversal initiation by exciting AVA/AVE neurons via electrical synapses mediated by the innexins UNC-7, UNC-9, and INX-1. This electrical connectivity may also function to maintain response strength. However, following RIM activation, glutamate released from RIM may then inhibit AVA/AVE and A-type motor neurons in the reversal circuit synaptically and/or extrasynaptically by turning on the inhibitory glutamate-gated chloride channel AVR-14, leading to the suppression of reversal frequency with time (Figure 5F). Indeed, in RIM-ablated worms, AVA/AVE neurons display a decrease in the amplitude of calcium spikes while exhibiting an increase in the frequency of calcium spikes, indicating that RIM neurons can both promote and suppress AVA/AVE activities. These findings also suggest that RIM promotes reversal initiation by potentiating the amplitude of calcium spikes in AVA/AVE neurons, but suppresses reversal probability by inhibiting the frequency of calcium spikes in these neurons, thereby providing a circuit mechanism underlying the dual-role of RIM in regulating reversal behavior.

Our data show that individual neurons in a neural circuit can regulate distinct features of a behavior by using either electrical or chemical transmission to communicate with other neurons in the circuit. Notably, these two modes of transmission are temporally distinct, as electrical transmission via gap junctions is rapid while chemical transmission occurs at a slower pace (Dong et al., 2018). This differential temporal pattern of information processing may explain why RIM initially promotes reversal initiation and subsequently supress reversal probability. Our findings suggest that the identified circuit is able to process temporal information to both promote and suppress motor output. Processing differential temporal patterns to fine tune circuit functions offers an excellent coding strategy for behavioral control.

In the locust, the dorsal uncrossed bundle (DUB) neurons and the lobula giant movement detector (LGMD) also form similar connections that process time-varying information (Wang et al., 2018), suggesting that similar mechanisms may operate in other species.

Complex brain functions are traditionally believed to depend on a vast number of neurons (Herculano-Houzel, 2012). However, increasing evidence suggests that it may also rely on multiple functions of single neurons. This phenomenon has been observed in both invertebrate and vertebrate brains (Briggman and Kristan, 2008; Li et al., 2014; Rigotti et al., 2013). The observation that RIM regulates multiple features of reversal behavior and also contributes to motor adaptation in *C. elegans* indicates that this neuron is multi-functional. In addition to RIM neurons, many other neurons are also multi-functional in *C. elegans*. For example, AIY interneurons are multi-functional and can both regulate reversals and adjust locomotion speed (Li et al., 2014). The AIB interneuron pair can regulate both locomotion and feeding behaviour (Zou et al., 2018). A single pair of PVD sensory neurons are able to regulate proprioception as well as responses to harsh touch (Tao et al., 2019). SMD neurons are also multifunctional and play roles at multiple hierarchical levels such as fast head casting and omega turn behaviors (Kaplan et al., 2020). This growing body of evidence indicates that complex brain functions rely on not only the vast number of neurons, but also multiple functions of individual neurons. Importantly, in the *C. elegans* connectome, many neurons form similar connection patterns like the circuit described here, indicating that the temporal coding strategy adopted by RIM to relay distinct information within a circuit could be widely employed in neural network integration and behavioral control.

## Methods

### Strains

*WT*: N2. TQ800: lite-1(xu7). TQ440: akIs3[Pnmr-1::gfp]. TQ3032: lite-1(xu7); xuEx1040[Pnmr-1::GCaMP3.0 + Pnmr-1::DsRed2b]. TQ6292: lite-1(xu7); xuEx2167[Pcex-1::GCaMP6f+Pgcy-13::sl2:mcherry]. TQ6744: lite-1(xu7); xuEx2257[Punc-4::tetx::sl2::yfp]. TQ6745: lite-1(xu7); xuEx2257[Punc-4::tetx::sl2::yfp]; xuEx1932[pgcy-13::TeTx-sl2-YFP;pgcy-13::dsRed2b; pnlp-12::dsRed2b]. TQ6875: xuEx1932[pgcy-13::TeTx-sl2-YFP;pgcy-13::dsRed2b; pnlp-12::dsRed2b];xuIs219[Podr-2b(3a)::yfp+Punc-122d::gfp]. TQ7103: xuEx2595[Pgcy-13::tdc-1 sense+antisense+Pgcy-13::DsRed]. TQ7119: xuEx1899[Punc-4::DsRed]; xuEx856[pBS-77::5’UTR + avr-14::sl2::yfp]. TQ7283: xuEx2693[Pgcy-13::Chrimson::sl2::yfp]. TQ7313; xuEx2717[Pgcy-13::avr-14(genomic+cDNA)::sl2::YFP];avr-14(1302). TQ7262: xuEx2675[Pnmr-1::avr-14(genomic+cDNA)::sl2::yfp;avr-14(ad1302). TQ7264: xuEx2677[Pnpr-4::avr-14(genomic+cDNA)::sl2::yfp; avr-14(ad1302). TQ7267: xuEx2680[Punc-4::avr-14(genomic+cDNA)::sl2::yfp; avr-14(ad1302). TQ7269: xuEx2682[Pacr-5::avr-14(genomic+cDNA)::sl2::yfp; avr-14(ad1302). TQ7271: xuEx2684[Pacr-2::avr-14(genomic+cDNA)::sl2::yfp; avr-14(ad1302). TQ7274: xuEx2687[Plgc-55::avr-14(genomic+cDNA)::sl2::yfp; avr-14(ad1302). TQ7280: xuEx2593[Pgcy-13::eat-4 sense+antisense+Pgcy-13::DsRed]. TQ7281: avr-14(ad1302)I; xuEx2593[Pgcy-13::eat-4 sense+antisense+Pgcy-13::DsRed]. TQ7324; xuEx2693[Pgcy-13::Chrimson::sl2::YFP]; inx-1(tm3524). TQ7326: xuEx2693[Pgcy13::chrimson::mcherry; unc-7(e5). TQ7325: xuEx2693[Pgcy13::chrimson::mcherry];unc-9(e101). TQ7327: xuEx2730[P gcy-13::tetx;Punc-122::GFP]; xuEx2693[Pgcy-13::chrimsom::mcherry]. TQ7365: xuEx2765[Pnmr-1:: avr-14(genomic+cDNA)::sl2::yfp;Punc-4:: avr-14(genomic+cDNA)::sl2::yfp]; avr-14(ad1302). TQ7332: xuEx2793[Pgcy-13::tetx::YFP;Punc-122::GFP];xuEx1040[Pnmr-1::GCaMP3.0 + Pnmr-1::DsRed2b]. TQ7399:xuEx2795[Pnmr-1::avr-14::GFP;Punc-4::avr-14::gFP];xuEx2766[Pgcy-13::eat-4 RNAi;Punc-122::GFP];;avr-14. TQ7340: xuEx2751[Pnpr-4::chrimson::mcherry];xuEx2793[Pgcy-13::tetx;Punc-122::GFP]. TQ7441: unc-9(e101);inx-1(tm3524);xuEx2693[Pgcy-13::Chrimson::sl2::yfp]. TQ7348: xuEx2751[Pnpr-4::chrimson::mcherry]. TQ7568: unc-7(e5);inx-1(tm3524); xuEx2693[Pgcy-13::chrimson::sl2::mcherry]. TQ7553: unc-7(e5);inx-1(tm3524);xuEx2751[Pnpr-4::chrimson::sl2::mcherry]. TQ7710: xuEx2887[Pnmr-1::sl2::mcherry2+Pinx-1L::sl2::YFP]. TQ7711: xuEx2888[Pnmr-1::sl2::mcherry2]+xuEx856[pBS-77::5’UTR + avr-14::sl2::yfp]. TQ8002: xuEx2316[pgcy-13::tdc-1(s+as)+pgcy-13::dsRed2b+pnlp-12::dsRed2b]. TQ8003: xuEx2323[pgcy-13::eat-4RNAi+pgcy-13::sl2::CFP+pnlp-12::dsRed2b]. TQ8004: xuEx2320[pgcy-13::unc-31(s+as)+pgcy-13::sl2-CFP+pnlp-12::dsRed2b].

### Laser ablation, optogenetics, and behavior

Laser ablations were performed on L1 or L2 worms (Bargmann and Avery, 1995). The transgene *Pnmr-1::gfp* was included in worms to help identify AVA, AVD, AVE and RIM. Control groups of animals underwent surgical preparation without laser irradiation.

Optogenetic interrogation of reversal initiation was performed as previously described (Piggott et al., 2011). Briefly, worms were grown on NGM plates supplied with 5 μM all-trans-retinal. Day 1 adult worms were tested on retinal-free NGM plates spread with a thin layer of OP50 bacteria. Amber light (5 s pulse; 590nm; ~0.2 mW/mm2) was delivered from a home-made LED light source to activate Chrimson to trigger behaviours. Animal behaviors were recorded and analyzed using the Wormlab system (MBF Bioscience). Each trial included five animals and at least five trials were performed for each group. Reversals were scored as positive responses if the animal stopped forward movement and initiated a reversal lasting at least half of one head swing upon light stimulation.

Spontaneous reversal frequency was analyzed using an automated worm tracking system as described previously (Feng et al., 2006; Li et al., 2006; Piggott et al., 2011). Day 1 adult worms were transferred to no food NGM plates for tracking and reversal frequency was recorded for 10-16 min.

### Calcium imaging

To elimiate intrinsic response to blue light that excites GCaMP (Liu et al., 2010; Ward et al., 2008), strains used for calcium imaging carried a mutation in *lite-1* gene that encodes a light-sensing receptor (Gong et al., 2016). Calcium imaging was performed on freely-behaving worms (Piggott et al., 2011). This system consists of an upright fluorescence stereomicroscope (Stemi SV11 M2 BIO), a dual-view beamsplitter, a X-Y motorized stage (Prior H101A) and an Andor EMCCD camera. The genetically-encoded calcium sensor GCaMPs (GCaMP 3.0 or GCaMP6f) were introduced into different neurons using neuron-specific promoters to observe calcium responses, and the red florescent protein DsRed was used as a reference channel for ratiometric imaging. A home-made software was used to coordinate the motorized stage and an Andor iXon EMCCD camera to track animal behaviours as well as capture GCaMP/DsRed signal. Day 1 adult worms were transferred to no food NGM plates for imaging. All experiments were conducted under the standard laboratory conditions (20°C, 30% humidity). Data processing was conducted using home-made software. GCaMP and DsRed ratio was calculated to indicate calcium responses.

#### Acknowledgments

Some strains were obtained from the CGC and Knockout Consortiums in the U.S.A. and Japan. E.A.R received pre-doctoral training grant support from the NIA and NIDCD. This work was supported by a grant from the NIGMS (to X.Z.S.X).

---

**Figure S1.**
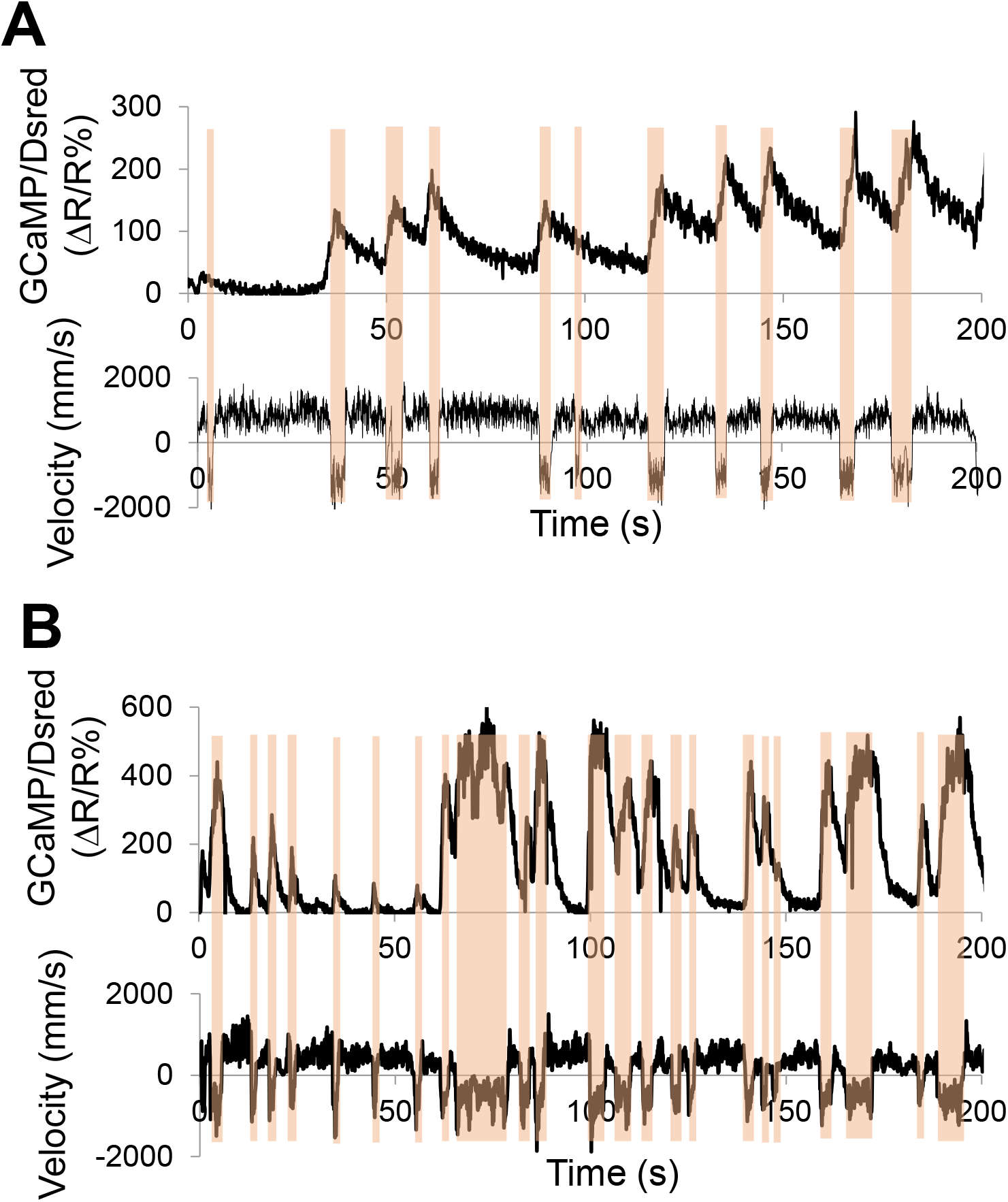
Additional calcium imaging traces. **(A-B)** Sample calcium traces of AVA/AVE neurons **(A)** and RIM neurons **(B)** in freely-moving animals. Calcium imaging was conducted with freely moving animals using the CARIBN system. Upper panels are calcium ratio traces. Lower panels are the velocity traces. Amber bars labeled reversal events. Most of the calcium spikes in AVA/AVE and RIM neurons are tightly coupled with reversals. The *nmr-1* promoter was used to express GCaMP3 transgene in AVA and AVE neurons, and the *cex-1* promoter was used to express GCaMP6f transgene in RIM neurons.

**Figure S2.**
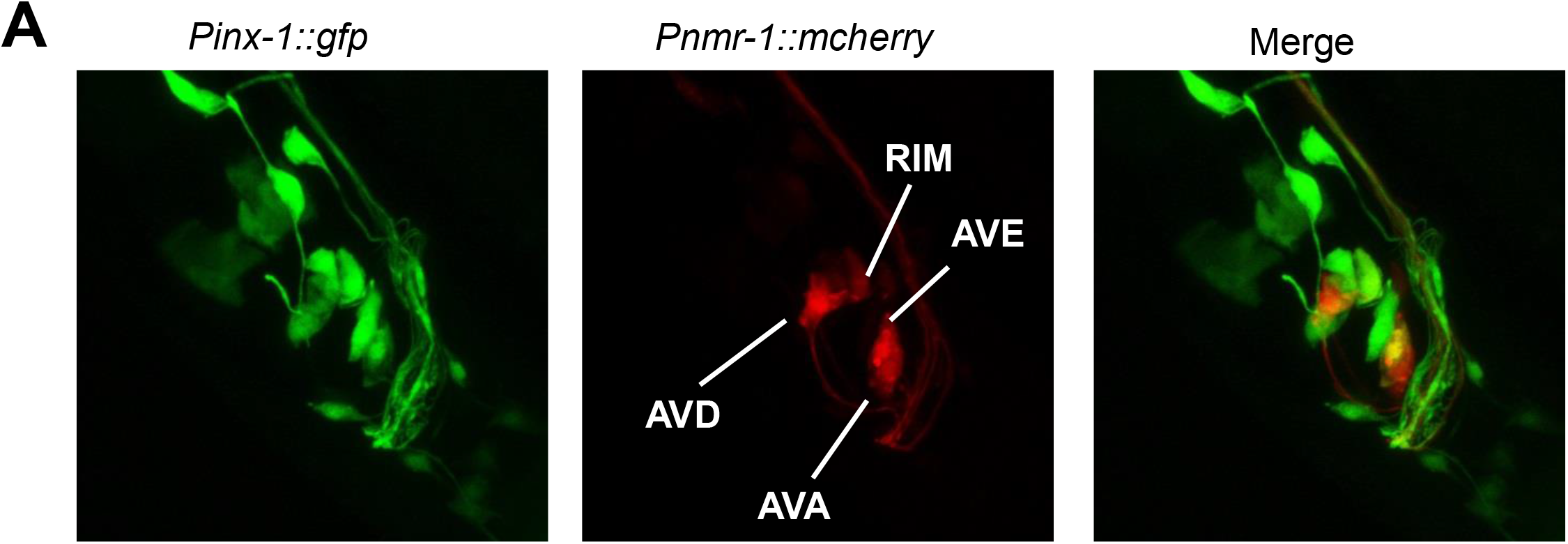
Expression pattern of *inx-1*. *Pnmr-1::mcherry* transgene labels the interneurons AVA, AVD, AVE and RIM in the head region. *Pinx-1::gfp* labels all of these neurons and other head neurons. *Pinx-1*: 3.2 kb promoter including 70bp coding region.

**Figure S3.**
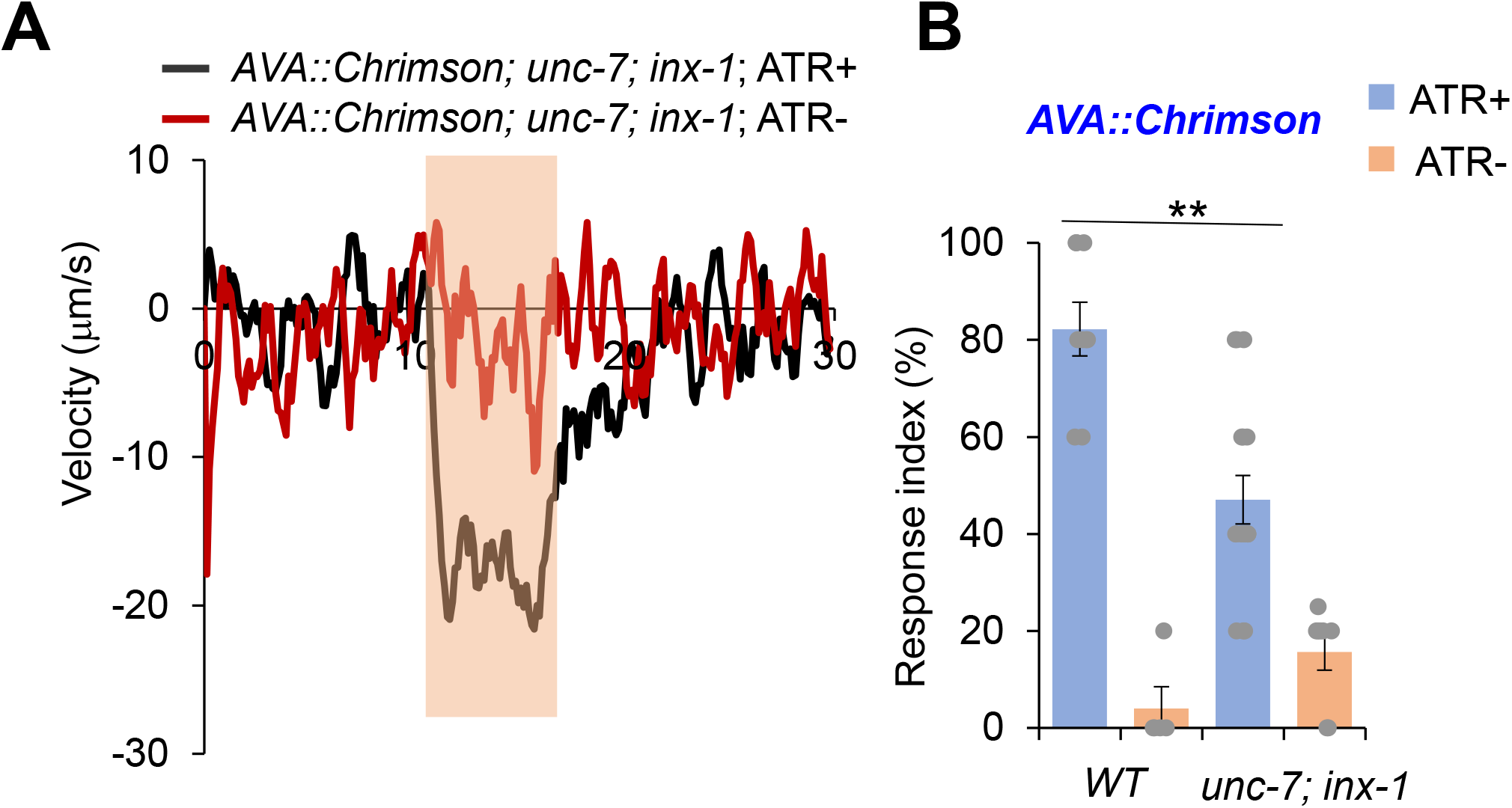
Activation of AVA neurons using optogenetics triggers reversals in *unc-7;inx-1* double mutant. **(A)** Reversals can still be triggered by AVA::Chrimson transgene in *unc-7;inx-1* mutants. Average traces. Bar in amber labels light stimulation segment. ATR: all-trans-retinal. **(B)** Bar graph shows reversal index quantification from (A). Error bars: SEM. n≥5. **p=5.525e-05 (ANOVA with Tukey’s HSD test).

**Figure S4.**
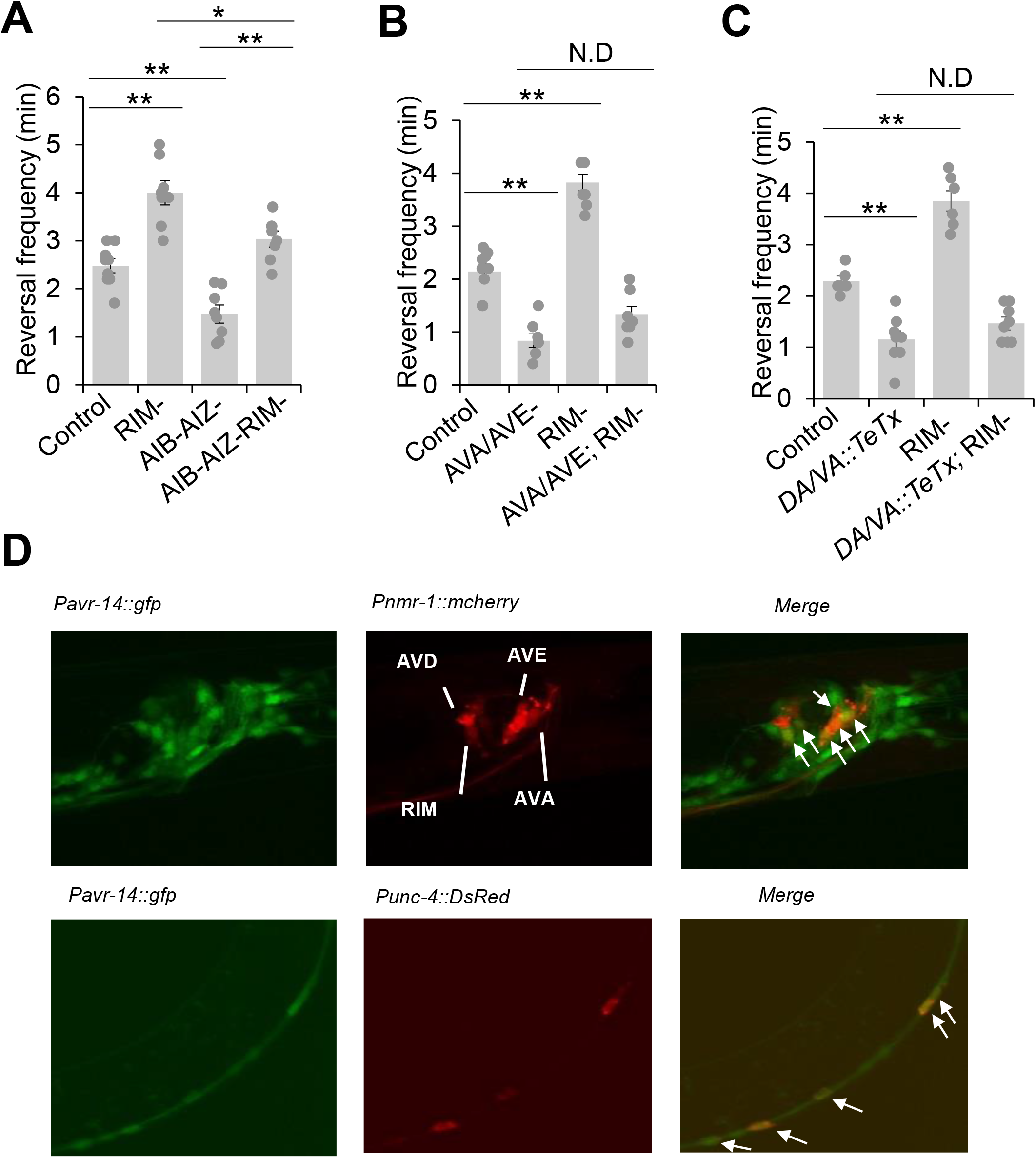
RIM suppresses reversal through AVA/AVE and A-type motor neurons and *avr-14*. **(A)** Ablation of the AIB and AIZ neurons does not the hyper-reversal phenotype in RIM-ablated worms. Bar graph shows quantification of reversal frequency. Error bars: SEM. n≥8. **p=1.252e-05 between control and RIM-, **p=0.002314 between control and AIB-AIZ-, *p=0.005383 between RIM- and AIB-AIZ-RIM-, **p=1.255e-05 (ANOVA with Tukey’s HSD test) between AIB-AIZ- and AIB-AIZ-RIM-. **(B)** Ablation of the AVA and AVE neurons largely suppresses the hyper-reversal phenotype in RIM-ablated worms. Bar graph shows quantification of reversal frequency. Error bars: SEM. n≥7. **p= 1.267e-05 between control and AVA/AVE-, **p= 1.126e-05 between control and RIM-, p= 0.1046 (ANOVA with Tukey’s HSD test) between AVA/AVE- and AVA/AVE-;RIM-. **(C)** Blocking the chemical transmission of A-type motor neurons using TeTx also suppresses the hyper-reversal phenotype in RIM-ablated worms. Bar graph shows quantification of reversal frequency. Error bars: SEM. n≥6. **p= 0.0005398 between control and DA/VA::Tetx, **p= 1.838e-05 between control and RIM-, p= 0.7303 (ANOVA with Tukey’s HSD test) between *DA/VA::TeTx* and *DA/VA::TeTx;* RIM-. **(D)** Expression pattern of *avr-14*. *Pnmr-1::mCherry* labels AVA, AVD, AVE and RIM neurons, and *Punc-4::DsRed* labels A-type motor neurons. White arrows indicate the overlapping neurons (RIM, AVA, AVE and A type motor neurons).

**Figure S5.**
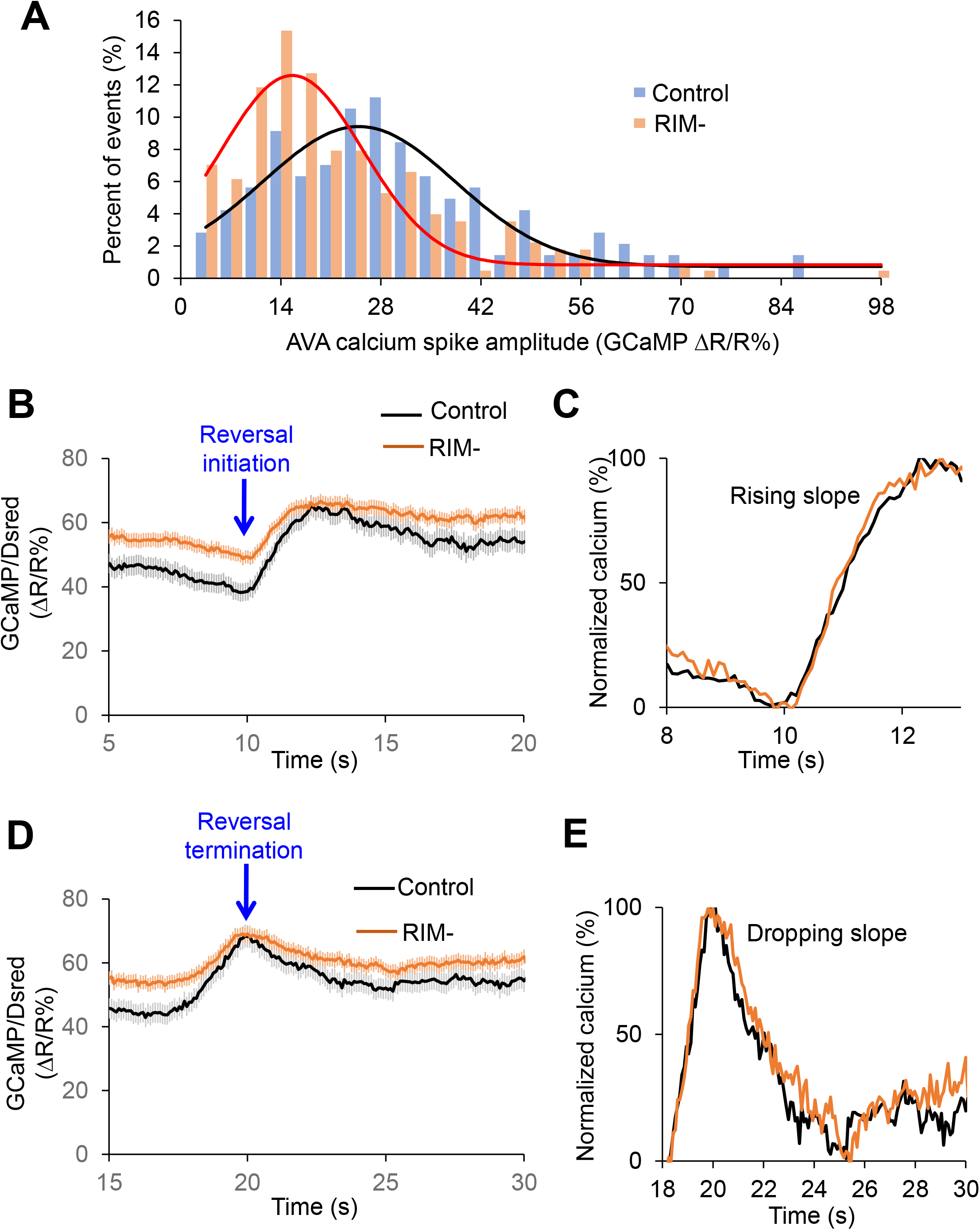
RIM ablation does not alter the kinetics of calcium spike in AVA/AVE neurons. **(A)** RIM ablation left-shifted the amplitude distribution pattern of calcium spikes in AVA/AVE neurons. Histograms are fitted with Gauss function. Only events with amplitude >0 are shown. Control: n=151; RIM-: n=264. **(B-C)** RIM ablation does not alter the rising slope of calcium spikes in AVA/AVE neurons. Arrow in (B) points to the time point of reversal initiation. (C) Normalized traces. **(D-E)** RIM ablation does not alter the dropping slope of calcium spikes in AVA/AVE neurons. Arrow in (D) points to the end of reversal (reversal termination). (E) Normalized traces.

## Notes

### Competing Interest Statement

The authors have declared no competing interest.

## Reference

Alkema, M.J., Hunter-Ensor, M., Ringstad, N., and Horvitz, H.R. (2005). Tyramine Functions independently of octopamine in the Caenorhabditis elegans nervous system. Neuron 46, 247–260.

Altun, Z.F., Chen, B., Wang, Z.W., and Hall, D.H. (2009). High resolution map of Caenorhabditis elegans gap junction proteins. Dev Dyn 238, 1936–1950.

Balaban, P.M. (2002). Cellular mechanisms of behavioral plasticity in terrestrial snail. Neurosci Biobehav Rev 26, 597–630.

Bargmann, C.I., and Avery, L. (1995). Laser killing of cells in Caenorhabditis elegans. Methods Cell Biol 48, 225–250.

Bhattacharya, A., Aghayeva, U., Berghoff, E.G., and Hobert, O. (2019). Plasticity of the Electrical Connectome of C. elegans. Cell 176, 1174–1189 e1116.

Briggman, K.L., and Kristan, W.B. (2008). Multifunctional pattern-generating circuits. Annu Rev Neurosci 31, 271–294.

Chalfie, M., Sulston, J.E., White, J.G., Southgate, E., Thomson, J.N., and Brenner, S. (1985). The neural circuit for touch sensitivity in Caenorhabditis elegans. J Neurosci 5, 956–964.

Chen, T.W., Wardill, T.J., Sun, Y., Pulver, S.R., Renninger, S.L., Baohan, A., Schreiter, E.R., Kerr, R.A., Orger, M.B., Jayaraman, V., et al. (2013). Ultrasensitive fluorescent proteins for imaging neuronal activity. Nature 499, 295–300.

Cote, M.P., Murray, L.M., and Knikou, M. (2018). Spinal Control of Locomotion: Individual Neurons, Their Circuits and Functions. Front Physiol 9, 784.

de Bono, M., and Maricq, A.V. (2005). Neuronal substrates of complex behaviors in C. elegans. Annu Rev Neurosci 28, 451–501.

Dent, J.A., Smith, M.M., Vassilatis, D.K., and Avery, L. (2000). The genetics of ivermectin resistance in Caenorhabditis elegans. Proc Natl Acad Sci U S A 97, 2674–2679.

Dong, A., Liu, S., and Li, Y. (2018). Gap Junctions in the Nervous System: Probing Functional Connections Using New Imaging Approaches. Front Cell Neurosci 12, 320.

Feng, Z., Li, W., Ward, A., Piggott, B.J., Larkspur, E.R., Sternberg, P.W., and Xu, X.Z. (2006). A C. elegans model of nicotine-dependent behavior: regulation by TRP-family channels. Cell 127, 621–633.

Gong, J., Yuan, Y., Ward, A., Kang, L., Zhang, B., Wu, Z., Peng, J., Feng, Z., Liu, J., and Xu, X.Z.S. (2016). The C. elegans Taste Receptor Homolog LITE-1 Is a Photoreceptor. Cell 167, 1252–1263 e1210.

Gray, J.M., Hill, J.J., and Bargmann, C.I. (2005). A circuit for navigation in Caenorhabditis elegans. Proc Natl Acad Sci U S A 102, 3184–3191.

Guo, Z.V., Hart, A.C., and Ramanathan, S. (2009). Optical interrogation of neural circuits in Caenorhabditis elegans. Nat Methods 6, 891–896.

Hart, A.C., and Chao, M.Y. (2010). From Odors to Behaviors in Caenorhabditis elegans. In The Neurobiology of Olfaction, A. Menini, ed.(Boca Raton (FL)).

Herculano-Houzel, S. (2012). The remarkable, yet not extraordinary, human brain as a scaled-up primate brain and its associated cost. Proc Natl Acad Sci U S A 109 Suppl 1, 10661–10668.

Hilliard, M.A., Bargmann, C.I., and Bazzicalupo, P. (2002). C. elegans responds to chemical repellents by integrating sensory inputs from the head and the tail. Curr Biol 12, 730–734.

Kaplan, H.S., Salazar Thula, O., Khoss, N., and Zimmer, M. (2020). Nested Neuronal Dynamics Orchestrate a Behavioral Hierarchy across Timescales. Neuron 105, 562–576 e569.

Kato, S., Kaplan, H.S., Schrodel, T., Skora, S., Lindsay, T.H., Yemini, E., Lockery, S., and Zimmer, M. (2015). Global brain dynamics embed the motor command sequence of Caenorhabditis elegans. Cell 163, 656–669.

Kawano, T., Po, M.D., Gao, S., Leung, G., Ryu, W.S., and Zhen, M. (2011). An imbalancing act: gap junctions reduce the backward motor circuit activity to bias C. elegans for forward locomotion. Neuron 72, 572–586.

Kim, K., and Li, C. (2004). Expression and regulation of an FMRFamide-related neuropeptide gene family in Caenorhabditis elegans. J Comp Neurol 475, 540–550.

Klapoetke, N.C., Murata, Y., Kim, S.S., Pulver, S.R., Birdsey-Benson, A., Cho, Y.K., Morimoto, T.K., Chuong, A.S., Carpenter, E.J., Tian, Z., et al. (2014). Independent optical excitation of distinct neural populations. Nat Methods 11, 338–346.

Lee, R.Y., Sawin, E.R., Chalfie, M., Horvitz, H.R., and Avery, L. (1999). EAT-4, a homolog of a mammalian sodium-dependent inorganic phosphate cotransporter, is necessary for glutamatergic neurotransmission in caenorhabditis elegans. J Neurosci 19, 159–167.

Li, W., Feng, Z., Sternberg, P.W., and Xu, X.Z. (2006). A C. elegans stretch receptor neuron revealed by a mechanosensitive TRP channel homologue. Nature 440, 684–687.

Li, Z.Y., Liu, J., Zheng, M.H., and Xu, X.Z.S. (2014). Encoding of Both Analog- and Digital-like Behavioral Outputs by One C. elegans Interneuron. Cell 159, 751–765.

Liu, J., Ward, A., Gao, J., Dong, Y., Nishio, N., Inada, H., Kang, L., Yu, Y., Ma, D., Xu, T., et al. (2010). C. elegans phototransduction requires a G protein-dependent cGMP pathway and a taste receptor homolog. Nat Neurosci 13, 715–722.

Pellizzari, R., Rossetto, O., Schiavo, G., and Montecucco, C. (1999). Tetanus and botulinum neurotoxins: mechanism of action and therapeutic uses. Philos Trans R Soc Lond B Biol Sci 354, 259–268.

Pierce-Shimomura, J.T., Chen, B.L., Mun, J.J., Ho, R., Sarkis, R., and McIntire, S.L. (2008). Genetic analysis of crawling and swimming locomotory patterns in C. elegans. Proc Natl Acad Sci U S A 105, 20982–20987.

Piggott, B.J., Liu, J., Feng, Z., Wescott, S.A., and Xu, X.Z. (2011). The neural circuits and synaptic mechanisms underlying motor initiation in C. elegans. Cell 147, 922–933.

Pirri, J.K., McPherson, A.D., Donnelly, J.L., Francis, M.M., and Alkema, M.J. (2009). A tyramine-gated chloride channel coordinates distinct motor programs of a Caenorhabditis elegans escape response. Neuron 62, 526–538.

Purves, D., Augustine, G.J., Fitzpatrck, D., Hall, W.C., LaMantia, A.S., McNAMARA, J.O., and White, L.E. (2008). Movement and its central control. In Neuroscience (Sunderland: Sinauer Associates, Inc.), pp. 397–541.

Rigotti, M., Barak, O., Warden, M.R., Wang, X.J., Daw, N.D., Miller, E.K., and Fusi, S. (2013). The importance of mixed selectivity in complex cognitive tasks. Nature 497, 585–590.

Serrano-Saiz, E., Poole, R.J., Felton, T., Zhang, F., De La Cruz, E.D., and Hobert, O. (2013). Modular control of glutamatergic neuronal identity in C. elegans by distinct homeodomain proteins. Cell 155, 659–673.

Speese, S., Petrie, M., Schuske, K., Ailion, M., Ann, K., Iwasaki, K., Jorgensen, E.M., and Martin, T.F. (2007). UNC-31 (CAPS) is required for dense-core vesicle but not synaptic vesicle exocytosis in Caenorhabditis elegans. J Neurosci 27, 6150–6162.

Tao, L., Porto, D., Li, Z., Fechner, S., Lee, S.A., Goodman, M.B., Xu, X.Z.S., Lu, H., and Shen, K. (2019). Parallel Processing of Two Mechanosensory Modalities by a Single Neuron in C. elegans. Dev Cell 51, 617–631 e613.

Wang, H., Dewell, R.B., Zhu, Y., and Gabbiani, F. (2018). Feedforward Inhibition Conveys Time-Varying Stimulus Information in a Collision Detection Circuit. Curr Biol 28, 1509–1521 e1503.

Ward, A., Liu, J., Feng, Z., and Xu, X.Z. (2008). Light-sensitive neurons and channels mediate phototaxis in C. elegans. Nat Neurosci 11, 916–922.

White, J.G., Southgate, E., Thomson, J.N., and Brenner, S. (1986). The structure of the nervous system of the nematode Caenorhabditis elegans. Philos Trans R Soc Lond B Biol Sci 314, 1–340.

Zheng, Y., Brockie, P.J., Mellem, J.E., Madsen, D.M., and Maricq, A.V. (1999). Neuronal control of locomotion in C. elegans is modified by a dominant mutation in the GLR-1 ionotropic glutamate receptor. Neuron 24, 347–361.

Zou, W., Fu, J., Zhang, H., Du, K., Huang, W., Yu, J., Li, S., Fan, Y., Baylis, H.A., Gao, S., et al. (2018). Decoding the intensity of sensory input by two glutamate receptors in one C. elegans interneuron. Nat Commun 9, 4311.

